# Proteasome activity is required for reovirus entry into cells

**DOI:** 10.1101/2023.05.10.540220

**Authors:** Andrew T. Abad, Andrew J. McNamara, Pranav Danthi

**Author notes:** To whom correspondence should be addressed: Department of Biology, Indiana University, Bloomington, IN 47405.

## Abstract

Since viruses have limited coding capacity in their genomes, they use host cell machinery to complete virtually every stage of their replication cycle. Mammalian orthoreovirus (reovirus) is comprised of two concentric protein shells, the inner core and the outer capsid. Following attachment to its receptor, reovirus enters the cell by receptor-mediated endocytosis. Within endosomes reovirus utilizes host acid-dependent proteases to process the viral outer capsid. Specifically, the outer capsid protein σ3 is degraded and μ1 is cleaved to form the disassembly intermediate infectious subvirion particles (ISVPs). ISVPs undergo additional conformational changes into ISVP*s that release small peptides which mediate the penetration of endosomal membranes.

Membrane penetration allows for delivery of the remaining viral core into the cytoplasm for subsequent gene expression. Here, we describe that the ubiquitin proteasome system controls an entry step of reovirus particles. We show that chemically inhibiting the proteasome blocks infection at a stage following ISVP formation but prior to transcriptional activation of cores. Specifically, inhibition of the proteasome prevents conformational changes in μ1 characteristic of ISVP-to-ISVP* conversion. In the absence of these conformational changes, cores are unable to be delivered and become transcriptionally active, thereby blocking viral replication. This work highlights a previously unknown way in which reovirus relies on host factors for successful replication.

**IMPORTANCE:** Due to their limited genetic capacity, viruses are reliant on multiple host systems to replicate successfully. Mammalian orthoreovirus (reovirus) is commonly used as a model system for understanding host-virus interactions. In this study, we identify the host ubiquitin proteasome system as a regulator of reovirus entry. Inhibition of the proteasome using a chemical inhibitor blocks reovirus uncoating. Blocking these events reduces subsequent replication of the virus. This work identifies that additional host factors controls reovirus entry.

## INTRODUCTION

Because most viruses have limited coding capacity in their genomes, they rely on cellular components to complete virtually every stage of their replication cycle. The reliance of viruses on host factors begins at the step of viral entry. During entry, viruses use host receptors for attachment, host proteins for trafficking to the appropriate cellular compartment and either specific host proteins or intracellular compartment with a distinct environment (such as pH, oxidative state) for uncoating or disassembly (1). Identification of such factors and an appreciation of the mechanism by which these factors promote virus replication can reveal previously unknown aspects of host cell biology and expose virus’ Achilles heel that can be exploited for the development and deployment of antiviral agents prior to a stage when genome replication begins.

Studies on the entry of mammalian orthoreovirus (reovirus), have been instrumental in revealing how multilayered, nonenveloped viruses complete the initial stages of infection to launch genome replication (2). Work on reovirus has also uncovered conserved aspects of the entry process of unrelated viruses. Particles of reovirus are 85 nm in diameter and are comprised of two concentric protein shells, designated as the outer capsid and the inner core (3, 4). The core encapsidates a 10-segmented dsRNA genome along with the RNA-dependent RNA polymerase and associated viral cofactors. During entry, an intact core particle is delivered into the cytoplasm for replication and gene expression. Cytoplasmic delivery of the core requires membrane disruption induced by viral outer capsid components (5). However, interaction of the particle with multiple host components is required prior to this event.

Reovirus particles initiate infection by attaching to host receptors. Following low affinity interaction with cell surface glycans, reovirus particles engage junctional adhesion molecule-A (JAM-A)(6–10). Receptor bound reovirus can be internalized into cells by endocytosis pathways dependent on either clathrin or caveolin or by macropinocytosis (11–15). Among these, clathrin-dependent uptake relies on interaction of the particle with β1 integrins (16). Once within a low pH compartment, the particle is acted upon by acid-dependent cathepsin proteases (17). These proteases, specifically cathepsin B and L in fibroblasts, remove the σ3 outer-capsid protein and cleave the μ1 outer capsid protein to generate an intermediate referred to as the infectious subvirion particle (ISVP). ISVPs undergo a conformational transition to form ISVP*s and concomitantly release μ1-derived peptides that help deliver viral cores into the cytoplasm (18, 19).

In this study, we identify a new function for the ubiquitin proteasome system for the early stages of entry. We demonstrate that treatment of cells with a proteasome inhibitor blocks infection following ISVP formation but prior to transcriptional activation of cores. Specifically, changes in the μ1 protein that occur during ISVP-to-ISVP* conversion are blocked by inhibition of the proteasome. The absence of structural transition prevents core delivery and transcriptional activation of incoming viral particles consequently diminishing viral replication.

## MATERIALS AND METHODS

### Cells

Spinner-adapted Murine L929 cells were maintained in Joklik’s MEM (Lonza) supplemented to contain 5% fetal bovine serum (FBS) (Invitrogen), 2 mM L-glutamine (Invitrogen), 100 U/ml penicillin (Invitrogen), 100 μg/ml streptomycin (Invitrogen), and 25 ng/ml amphotericin B (Sigma-Aldrich). HeLa cells obtained from Melanie Marketon’s laboratory and maintained in high glucose DMEM (Corning) supplemented to contain 10% FBS (Invitrogen), 2 mM L-glutamine (Invitrogen), 100 U/ml penicillin (Invitrogen), 100 μg/ml streptomycin (Invitrogen).

### Reagents and antibodies

MG132, PSI and Epoxomicin were purchased from Cayman Chemicals, Millipore, and Sigma-Aldrich respectively. Ribavirin was purchased from Sigma-Aldrich. Each inhibitor was resuspended in DMSO (Sigma-Aldrich). Anti-reovirus polyclonal antisera were generated by immunizing rabbits with UV-inactivated mixture of purified virions of T1L and T3D. Anti-core antisera were obtained from the Maya Shmulevitz laboratory (University of Alberta)(20). mAb 8H6 and 4A3, previously described, were obtained from the Terry Dermody laboratory (University of Pittsburgh)(21). Antisera specific to PSTAIR were obtained from Sigma-Aldrich.

### Virus purification

Reovirus strain T3D^C^ was obtained from John Parker. Strain T1L were regenerated by plasmid based reverse genetics (22). Purified reovirus virions were generated using second- or third-passage L-cell lysates stocks of reovirus. Viral particles were Vertrel-XF (Dupont) extracted from infected cell lysates, layered onto 1.2-to 1.4-g/cm^3^ CsCl gradients, and centrifuged at 187,183 x *g* for 4 h. Bands corresponding to virions (1.36 g/cm^3^) were collected and dialyzed in virion-storage buffer (150 mM NaCl, 15 mM MgCl_2_, 10 mM Tris-HCl [pH 7.4]) (23).

### Generation of ISVPs in vitro

ISVPs were generated by incubation of 2 x 10^12^ virions with 200 µg/ml of CHT in a total volume of 0.1 ml at 32°C in virion storage buffer for 20 min (24). Proteolysis was terminated by addition of 2 mM phenylmethylsulphonyl fluoride (PMSF) and incubation of reactions on ice.

### Plaque assays

Plaque assays to determine infectivity were performed as previously described with some modifications (23, 25). Briefly, control or heat-treated virus samples were diluted into PBS supplemented with 2 mM MgCl_2_ (PBS^Mg^). L cell monolayers in 6-well plates (Greiner Bio-One) were infected with 100 μl of diluted virus for 1 h at room temperature. Following the viral attachment incubation, the monolayers were overlaid with 4 ml of serum-free medium 199 (Sigma-Aldrich) supplemented with 1% Bacto Agar (BD Biosciences), 10 μg/ml TLCK-treated chymotrypsin (Worthington, Biochemical), 2 mM L-glutamine (Invitrogen), 100 U/ml penicillin (Invitrogen), 100 μg/ml streptomycin (Invitrogen), and 25 ng/ml amphotericin B (Sigma-Aldrich). The infected cells were incubated at 37°C, and plaques were counted 5 d post infection.

### Assessment of infectivity by indirect immunofluorescence

Monolayers of cells (4 x 10^4^) in clear bottom tissue culture treated 96-well plates (Corning) were washed with PBS and adsorbed with virions or ISVPs of the indicated reovirus strain at 4°C for 1 h.

Cells were incubated in medium supplemented with the appropriate inhibitor. Monolayers were fixed with methanol at -20°C for a minimum of 30 min and washed with PBS containing 0.5% Tween-20 (DPBS-T). Cells were then incubated with polyclonal rabbit anti-reovirus serum at a 1:1000 dilution in PBS with 1% BSA (DPBS-BSA) at 37°C for 60 min. Monolayers were washed twice with DPBS-T and incubated with DPBS-BSA for 37°C for 60 min followed by two washes with DPBS-T. Cells were stained with 1:1000 dilution of LI-COR CW 800 anti-rabbit immunoglobin G, 1:1000 dilution of Sapphire 700 (LI-COR) and DRAQ5 (Cell signaling Technology) at a concentration of 1:10000 for 37°C for 60 min. Monolayers were washed thrice with DPBS-T and fluorescence intensity was measured using the Odyssey Imaging System and the Image Studio Lite software (LI-COR). For each well, the ratio of fluorescence at 800 nm (for infected cells) and 700 nm (for total cells) was quantified. Infectivity in arbitrary units was quantified using the following formula. Infectivity = (Green/Red)_infected_-(Green/Red)_uninfected_ (26).

### Cell lysate preparation and immunoblotting

For preparation of whole cell lysates, cells were washed in phosphate-buffered saline (PBS) and lysed with 1X RIPA (50 mM Tris [pH 7.5], 50 mM NaCl, 1% TX-100, 1% DOC, 0.1% SDS, and 1 mM EDTA) containing a protease inhibitor cocktail (Roche) and 2 mM PMSF. The samples were centrifuged at 15000 × *g* for 10 min to remove debris. Cell lysates were resolved by electrophoresis on 10% polyacrylamide gels and transferred to nitrocellulose membranes. Membranes were blocked for at least 1 h in TBS containing T20 Starting Block (Thermo Fisher) and incubated with antisera against reovirus (1:5000) or PSTAIR (1:10000) at 4°C overnight. Membranes were washed three times for 5 min each with washing buffer (TBS containing 0.1% Tween-20) and incubated with 1:20000 dilution of Alexa Fluor conjugated goat anti-rabbit IgG (for reovirus polyclonal) or goat anti-mouse IgG (for PSTAIR) in blocking buffer. Following three washes, membranes were scanned using an Odyssey Infrared Imager (LI-COR).

### RT-qPCR

RNA was extracted from infected cells at various time intervals after infection using Total RNA mini kit (Biorad). RNA present in in vitro transcription reactions was directly subjected to reverse transcription without purification. For RT-qPCR, RNA was reverse transcribed with the High Capacity cDNA Reverse Transcription Kit (Applied Biosystems) using random hexamers. A 1:10 dilution of the cDNA was subjected to qPCR using SYBR Select Master Mix (Applied Biosystems). Fold increase in gene expression with respect to control samples (indicated in each figure legend) was measured using the ΔΔC_T_ method (27). Calculations for determining ΔΔC_T_ values and relative levels of gene expression were performed as follows: (i) fold increase in viral _gene expression = 2_ -[(T3D S1 CT – GAPDH CT)MG132 – (T3D S1 CT – GAPDH CT)DMSO_. Primer_ sequences are available upon request.

### In vitro transcription

ISVPs (2 x 10^12^ particles/ml) were incubated with 300 mM CsCl at 32°C for 20 min (19). 8 x 10^9^ Cs treated ISVPs in 20 μl reactions were incubated at 45°C for 2 h in a buffer containing 100 mM HEPES-KOH (pH 8.1), 10 mM MgCl2, 0.5 mM EDTA, 3.3 mM phosphoenol pyruvate, 100 μg/ml pyruvate kinase, 0.1 mM ATP, 0.1 mM CTP, 0.1 mM UTP, and 0.4 mM GTP. Negative control reactions contained no NTPs. DMSO or MG132 (5 μM) were added from a 10X stock. RNA accumulated was reverse transcribed using High Capacity cDNA Reverse Transcription Kit. A 1:10 dilution of the cDNA was subjected to qPCR using SYBR Select Master Mix (Applied Biosystems). Fold increase in gene expression with respect to control samples (indicated in each figure legend) was measured. Fold increase in viral gene expression = 2 ^-[(T1L S2 CT experimental – (T1L S2 CT control)^. Primer sequences are available upon request.

### Evaluation of reovirus uncoating

Monolayers of HeLa cells were grown on 8-well chambered slides (Falcon) at 37°C to ∼80% confluence. Cells were pretreated with 100 μg/mL cycloheximide (CHX) in DMEM with DMSO or MG132 (10 μM) for 1 h at 37°C. Cells were then inoculated with 10^5^ T3D^C^ ISVPs/cell for 1 h at 4°C with occasional rocking. Following incubation, the inoculum was removed and complete medium with 100 μg/mL CHX and DMSO or MG132 was added. Cells were then incubated for 2 h at 37°C. Cells were washed with PBS and fixed with 3.7% formaldehyde for at least 10 minutes at room temperature. Fixed cells were washed with PBS three times and permeabilized with PBS containing 0.1% Triton X-100 for 10 min at room temperature. After incubation, cells were washed with PBS three times and blocked with PBS containing 2.5% BSA for 30 min at room temperature. The blocking solution was then removed and cells were incubated with anti-core antibody and 4A3 (1:1000) or 8H6 (1:1000) in PBS containing 0.1% Triton X-100 and 0.1% BSA (wash solution) for 1 h at room temperature. Following incubation, cells were washed three times with wash solution. Cells were then incubated with goat anti-mouse Alexa Fluor 488 (1:1000, Invitrogen) and donkey anti-rabbit Alex Fluor 594 (1:1000, Life Technology) in wash solution for 1 h at room temperature in the dark. Following incubation, cells were washed twice with wash solution. Cells were washed a third time with wash solution containing 1 μg/mL DAPI for 10 min at room temperature in the dark. A coverslip was mounted with AquaPoly mounting solution (Polysciences). Confocal images were acquired using a Leica SP8 scanning confocal microscope controlled by Leica LAS-X software within the Indiana University Light Microscopy Imaging Center. Images were obtained using 63× oil-immersion objective and the white light laser, 405 nm laser, and HyD detectors. Three-dimensional image stacks were acquired by recording sequential sections through the *z*-axis. Images were processed with ImageJ (two-dimensional maximum-intensity projections)(28).

### α-sarcin co-entry assay

Monolayers of HeLa cells were grown in a 96-well black plate at 37°C to ∼90% confluency. Cells were pretreated with DMSO or MG132 (5 μM) for 1 h at 37°C. Following pretreatment, T3D^C^ or T1L ISVPs were adsorbed at an MOI of 10^5^ particles/cell for 1 h at 4°C with occasional rocking. The inoculum was then removed and complete media was added with DMSO or 5 μM MG132 plus 50 μg/mL α-sarcin. The cells were incubated for 2 h at 37°C. Puromycin was added to cells for a final concentration of 20 μg/mL for 30 min prior to fixation. At 2 h post infection, cells were washed with PBS and fixed with ice-cold methanol at -20°C. Fixed cells were probed for puromycin incorporation using a puromycin-specific antibody (1:1000, MABE343, EMD Millipore). The amount of puromycin incorporated was quantified using the method described above for infectivity measurement using LI-COR Odyssey.

## RESULTS

### Proteasome inhibition blocks multiple stages of reovirus infection

In the course of experiments to examine the turnover of host proteins in reovirus infected cells, we observed that treatment of cells at the start of infection with a proteasome inhibitor, MG132, diminished infection (data not shown)(29, 30). To gain a better understanding of the observations described above, we treated cultured cells with increasing concentration of MG132, a proteasome inhibitor that is widely used in the laboratory (30). We found that MG132 treatment of uninfected HeLa cells at doses up to 10 μM, did not significantly compromise cell viability over 18 h (data not shown). Based on these results, we used 5 μM MG132 for our experiments to test the effect of proteasome inhibition on reovirus infection. Unlike BV6, a known apoptosis inducer, 5 μM MG132 has minimal impact on cell viability in the time frame needed to analyze infection by virions or ISVPs (Figure 1A) (31). Analysis of infection efficiency by indirect immunofluorescence indicated that type 3 reovirus strain T3D^C^ failed to establish infection in cells treated with MG132 (Figure 1B). Consistent with this, we found that in comparison to the control (DMSO), MG132 treatment resulted in a ∼ 2.5 log_10_ reduction in virus output. Thus, inhibition of the proteasome potently blocks infection and replication by T3D^C^ (Figure 1C).

**Figure 1.**
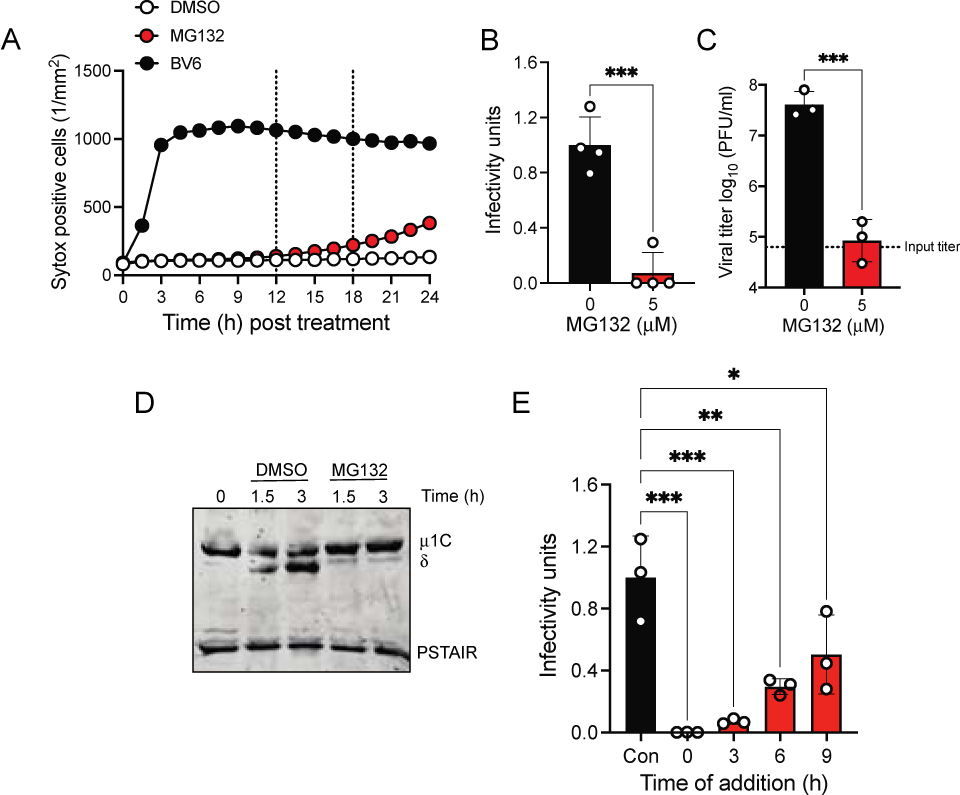
MG132 blocks reovirus infection (A) HeLa cells were incubated with DMSO, MG132 (5 μM), or BV6 (10 μM). The media was supplemented with 50 nM Sytox Green. Loss of cell viability, as measured by appearance of green fluorescence was monitored and quantified for a period of 24 h using Incucyte S3 imager and its associated software. Vertical lines indicate time points at which infection with virions (18 h) and ISVPs (12 h) were stopped in subsequent experiments. (B) HeLa cells were adsorbed with virions of T3D^C^ strain at MOI of 10 PFU/cell. After incubation at 37°C for 18 h in the presence of DMSO or MG132 (5 μM), the cells were subjected to indirect immunofluorescence assay using a LI-COR Odyssey scanner. Infectivity was determined by calculating intensity ratios at 800 nm (green fluorescence) representing viral antigen and 700 nm (red fluorescence) representing the cell monolayer. The 800/700 intensity ratio in absence of MG132 was set to 1. Infectivity for each independent infection and the sample mean are shown. Error bars indicate SD. ****, P < 0.001 as determined by Student’s t-test. (C) HeLa cells were infected with virions of T3D^C^ strain at MOI of 1 PFU/cell. Following incubation for 0 h (input titer) or 18 h in the presence of DMSO or MG132 (5 μM), viral titer was determined by plaque assay. Titer for each individual sample and sample mean are shown. ***, P < 0.005 as determined by Student’s t-test. (D) HeLa cells were adsorbed with virions of T3D^C^ strain at MOI of 10^4^ particles/cell. Following incubation for the indicated time intervals in the presence of ribavirin (200 μM) and either DMSO or MG132 (5 μM), the cells were lysed and subjected to immunoblotting using polyclonal sera against reovirus or PSTAIR. (E) HeLa cells were adsorbed with virions of T3D^C^ strain at MOI of 10 PFU/cell. Following 18 h incubation in the presence of DMSO (Con), or MG132 (5 μM) added at the indicated time intervals, the cells were subjected to indirect immunofluorescence assay using a LI-COR Odyssey scanner. Infectivity was determined by calculating intensity ratios at 800 nm (green fluorescence) representing viral antigen and 700 nm (red fluorescence) representing the cell monolayer. The 800/700 intensity ratio in absence of MG132 was set to 1. Infectivity for each independent infection and the sample mean are shown. Error bars indicate SD. *, P < 0.05; **, P< 0.01; ***, P < 0.005 as determined by one way ANOVA with Dunnett’s multiple comparison test.

Following attachment to receptors, reovirus particles are internalized and disassembled within the endosomal compartment by acid-dependent cathepsin B and/or L proteases to generate infectious subvirion particles (ISVPs)(17). Preventing ISVP formation by blocking endosomal acidification or by inhibiting the activity of cathepsins, results in blockade of infection (17, 32). However, MG132 is a protease inhibitor. In addition to the proteasome, it can also inhibit the activity of cathepsin proteases (33). Therefore, one way in which MG132 may impact infection is by preventing disassembly of the virus to generate ISVPs. To test if MG132 affects virus disassembly, we monitored the cleavage of μ1 outer-capsid protein from incoming virions as a hallmark of ISVP formation. As expected, μ1 is cleaved to δ in DMSO treated cells. In contrast, treatment of cells with MG132 blocked the generation of δ. These data indicate that MG132 blocks reovirus infection by preventing viral disassembly (Figure 1D).

Internalized reovirus particles disassemble within 2-3 h. Consequently, following this interval, infection is no longer susceptible to inhibition of cathepsin proteases (32, 34–36). To test if MG132 also functions in a similar time frame, we performed a time of addition experiment. To this end, we added MG132 to the media of cells at various times following infection and quantified infection by indirect immunofluorescence at 18 h following infection. Addition of MG132 at the start of infection (0 h) and 3 h post infection most potently blocked infection (Figure 1E). Interestingly, addition of MG132 at 6 h and 9 h following infection also inhibited infection. Though the effect of MG132 addition later in infection was less pronounced, infection was still significantly inhibited. These data raised the possibility that stages following ISVP formation are also impacted by inhibition of the proteasome.

### Proteasome inhibition blocks reovirus infection following proteolytic disassembly

ISVPs can be generated in vitro by treatment of virions with chymotrypsin. ISVPs generated in vitro can launch infection without the need for uptake into cathepsin-containing compartments and additional proteolytic processing (12, 17, 37). Experimental infection with ISVPs can therefore be used to define if a chemical or genetic perturbation impacts steps after disassembly. We infected cells with ISVPs and measured infection using indirect immunofluorescence at 12 h following infection. An earlier time point was selected because ISVPs complete their replication cycle within a shorter time period than virions. We found that infection of HeLa cells with in vitro generated ISVPs was also inhibited by MG132 (Figure 2A). Infection initiated by ISVPs in presence of proteasome inhibitors also led to a diminishment in infectious viral production (Figure 2B). We also confirmed that MG132 inhibited infection by ISVPs derived from type 1 reovirus strain, T1L (Figure 2C). Two other proteasome inhibitors, PSI and epoxomicin that are chemically distinct (33), also blocked infection by ISVP (Figure 2D). Together our data provide strong evidence that proteasome activity plays a virus serotype-independent role in the infectious cycle of reovirus.

**Figure 2.**
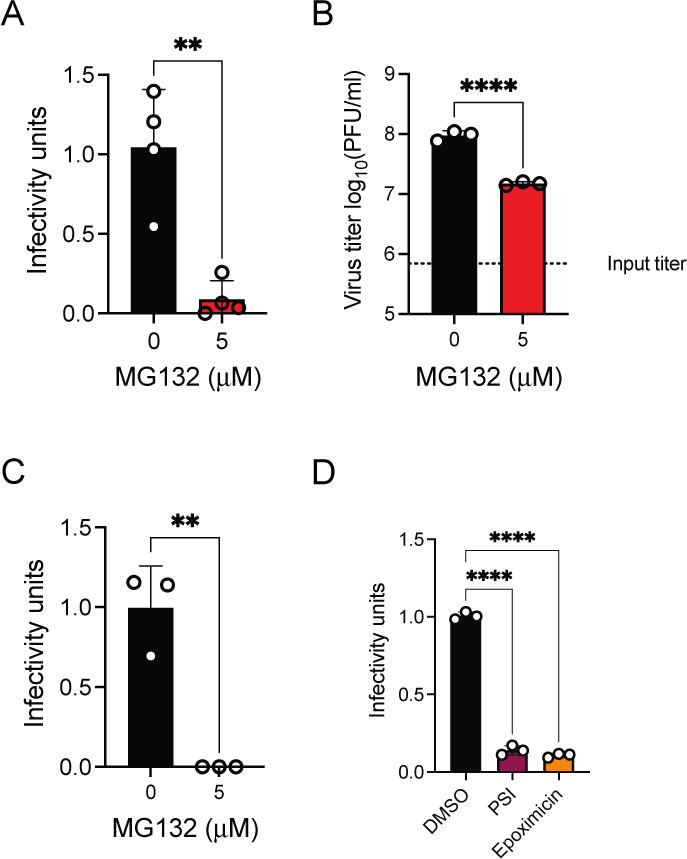
Proteasome inhibitors block infection by ISVPs. (A) HeLa cells were adsorbed with ISVPs of T3D^C^ strain at MOI of 10 PFU/cell. After incubation at 37°C for 12 h in the presence of DMSO or MG132 (5 μM), the cells were subjected to indirect immunofluorescence assay using a LI-COR Odyssey scanner. Infectivity was determined by calculating intensity ratios at 800 nm (green fluorescence) representing viral antigen and 700 nm (red fluorescence) representing the cell monolayer. The 800/700 intensity ratio in absence of MG132 was set to 1. Infectivity for each independent infection and the sample mean are shown. Error bars indicate SD. **, P < 0.01 as determined by Student’s t-test. (B) HeLa cells were infected with ISVPs of T3D^C^ strain at MOI of 1 PFU/cell. Following incubation for 0 h (input titer) or 12 h in the presence of DMSO or MG132 (5 μM), viral titer was determined by plaque assay. Titer for each individual sample and sample mean are shown. Error bars indicate SD. ****, P < 0.001 as determined by Student’s t-test. (C) HeLa cells were adsorbed with ISVPs of T1L strain at MOI of 10 PFU/cell. After incubation at 37°C for 12 h in the presence of DMSO or MG132 (5 μM), the cells were subjected to indirect immunofluorescence assay using a LI-COR Odyssey scanner. Infectivity was determined by calculating intensity ratios at 800 nm (green fluorescence) representing viral antigen and 700 nm (red fluorescence) representing the cell monolayer. The 800/700 intensity ratio in absence of MG132 was set to 1. Infectivity for each independent infection and the sample mean are shown. Error bars indicate SD. ***, P < 0.005 as determined by Student’s t-test. (D) HeLa cells were adsorbed with ISVPs of T3D^C^ strain at MOI of 10 PFU/cell. After incubation at 37°C for 12 h in the presence of DMSO, PSI (5 μM) or epoxomicin (400 nM), the cells were subjected to indirect immunofluorescence assay using a LI-COR Odyssey scanner. Infectivity was determined by calculating intensity ratios at 800 nm (green fluorescence) representing viral antigen and 700 nm (red fluorescence) representing the cell monolayer. The 800/700 intensity ratio in absence of MG132 was set to 1. Infectivity for each independent infection and the sample mean are shown. Error bars indicate SD. Error bars indicate SD. ****, P < 0.001 as determined by using one way ANOVA with Dunnett’s multiple comparison test.

### Steps subsequent to reovirus core delivery are not impacted by proteasome inhibition

Toward the goal of pinpointing the step at which MG132 blocks infection by ISVPs, we sought to define the time interval when infection with ISVPs is no longer sensitive to MG132. For these experiments, infection was initiated using ISVPs and MG132 was added to the media of the cells at different time intervals following infection. Consistent with our experiments above, addition of MG132 at the start of infection reduced infectivity (Figure 3A). The infection remained sensitive to MG132 when the drug was added at 2 h. However, addition of MG132 at 4 or 6 h post infection did not significantly impact infection. These data suggest that events during infection that occur soon after ISVP formation are dependent on the function of the proteasome. Such steps could include delivery of the core into the cytoplasm or the transcriptional activity of cores.

**Figure 3.**
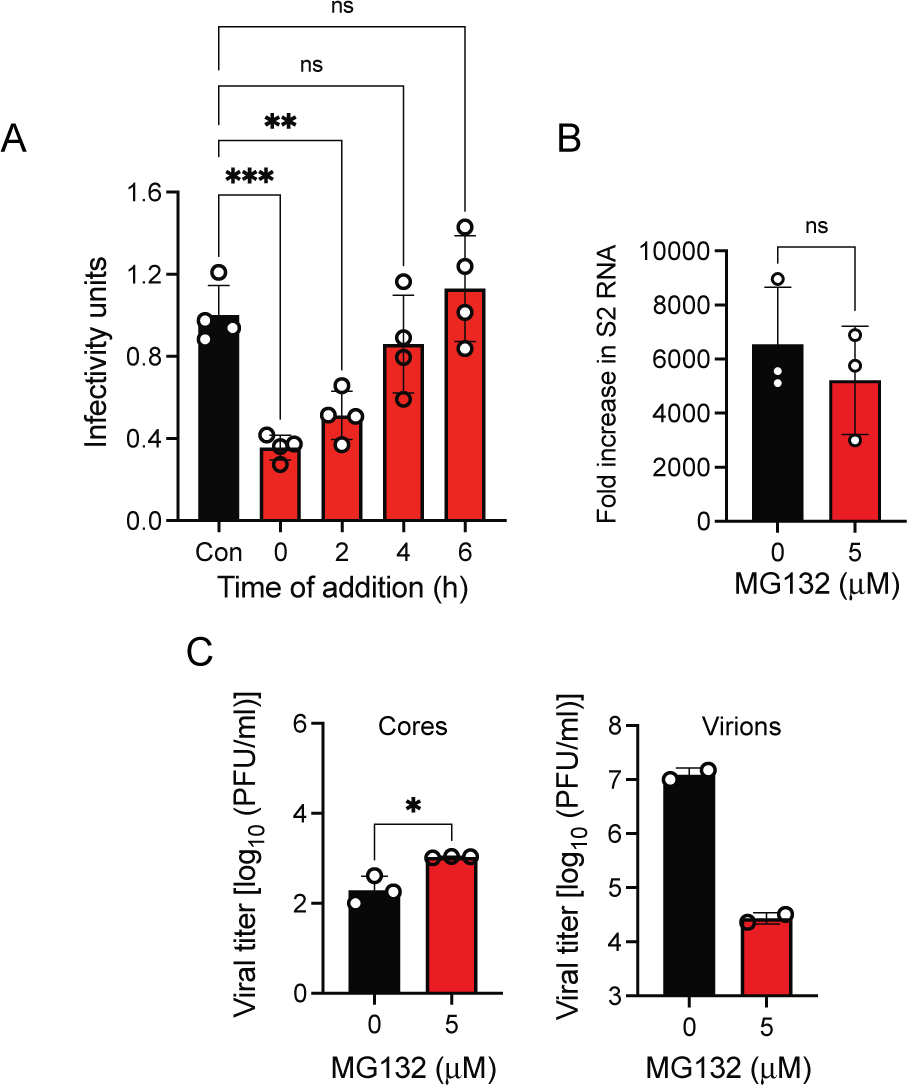
Proteasome inhibitor inhibits infection prior to core transcription. (A) HeLa cells were adsorbed with ISVPs of T3D^C^ strain at MOI of 10 PFU/cell. Following 12 h incubation in the presence of DMSO (Con), or MG132 (5 μM) added at the indicated time intervals, the cells were subjected to indirect immunofluorescence assay using a LI-COR Odyssey scanner. Infectivity was determined by calculating intensity ratios at 800 nm (green fluorescence) representing viral antigen and 700 nm (red fluorescence) representing the cell monolayer. The 800/700 intensity ratio in absence of MG132 was set to 1. Infectivity for each independent infection and the sample mean are shown. Error bars indicate SD. Error bars indicate SD. ****, P < 0.001 as determined by one way ANOVA with Dunnett’s multiple comparison test. (B) Reovirus ISVPs were incubated with CsCl and were subjected to in vitro transcription for 2 h in the presence of DMSO or MG132 (5 μM). Levels of accumulated T1L S2 mRNA was quantified by qPCR and comparative CT analysis. The amount of T1L S2 RNA detected in negative control transcription reactions was set to 1. Mean values for three independent in vitro transcriptions are shown. Error bars indicate SD. P values were determined by student’s t-test. ****, p < 0.001. (C) HeLa cells incubated in media containing DMSO or MG132 (5 μM) for 2 h were transfected with 10^4^ particles/cell of CsCl purified core particles of strain T1L using lipofectamine 2000. 24 h following transfection, viral titer was determined by plaque assay. Titer for each individual sample and sample mean are shown. Error bars indicate SD. Error bars indicate SD. ****, P < 0.001 as determined by Student’s t-test. In parallel experiments, HeLa cells were infected with virions of T3D^C^ strain at MOI of 1 PFU/cell. Following incubation 18 h in the presence of DMSO or MG132 (5 μM), viral titer was determined by plaque assay. ***, P < 0.005 as determined by Student’s t-test.

To measure transcriptional activity of reovirus particles in the presence of MG132, we used in vitro transcription. Transcriptase activity of in vitro generated ISVPs was stimulated by addition of CsCl (19). The RNA accumulated following 2 h of transcription was quantified by RT-qPCR using S2 gene specific primers. In vitro transcription from cores occurs with equivalent efficiency in the presence of DMSO or MG132 (Figure 3B). Thus, MG132 does not appear to directly impact the transcriptional function of reovirus particles via an off-target effect.

Since our in vitro transcription reactions do not contain any cellular proteins, it remains possible that, even though in vitro transcription is not affected by MG132, transcription from cores in cells is inhibited by MG132. To rule out this possibility, we exploited the capacity of transfected cores to launch infection. Cores lack the outer-capsid apparatus needed to attach and to enter cells. Therefore, they are not infectious when simply added to cells (38). However, they can launch infection upon transfection into cells (39). To test whether proteasome inhibition impacts recovery of infectious virus following core transfection, DMSO or MG132 treated cells were transfected with cores. Viral titer was determined at 24 h following transfection. This time point was selected to ensure that only output directly generated from core transfection and not subsequent spread of newly generated virus to neighboring cells was quantified. Consistent with expectations, cores added to cells without transfection reagent yielded no detectable progeny (data not shown). Transfection of cores into DMSO or MG132 treated cells allowed recovery of virus (Figure 3C). Importantly, the amount of infectious virus recovered was equivalent. Parallel infection of cells with virions in the presence of DMSO and MG132 confirmed that MG132 was effective at the concentration used in blocking virus replication (consistent with Figure 1). Because transfected cores remained capable of launching infection, these data indicate that proteasome activity is dispensable for reovirus infection once cores devoid of any residual outer capsid protein are delivered into the host cell.

### Blockade of proteasome activity reduces uncoating of ISVPs

Since the transcriptional activity of cores was not affected by MG132, we next turned to define whether core particles are successfully delivered into the cytoplasm. The core proteins are masked by μ1 fragments in ISVPs (4, 40). When ISVPs uncoat and deliver cores into the cytoplasm, μ1 fragments are released and the core becomes exposed (18). To determine whether these events are impacted by MG132, we evaluated the detectability of cores 2 h following infection with ISVPs. A mAb specific to the μ1 δ fragment, 8H6, was simultaneously used to confirm that ISVP internalization was not impacted. We observed that ISVPs marked by 8H6 were present within the cell following infection in presence of both DMSO and MG132 (Figure 4). Moreover, ISVP staining was visible in nearly every cell under both conditions. In contrast, DMSO treated cells showed significantly more core staining compared to cells treated with MG132. Intense core staining was found in significantly fewer cells following MG132 treatment. These data indicate that MG132 impacts delivery of cores into the cytoplasm.

**Figure 4.**
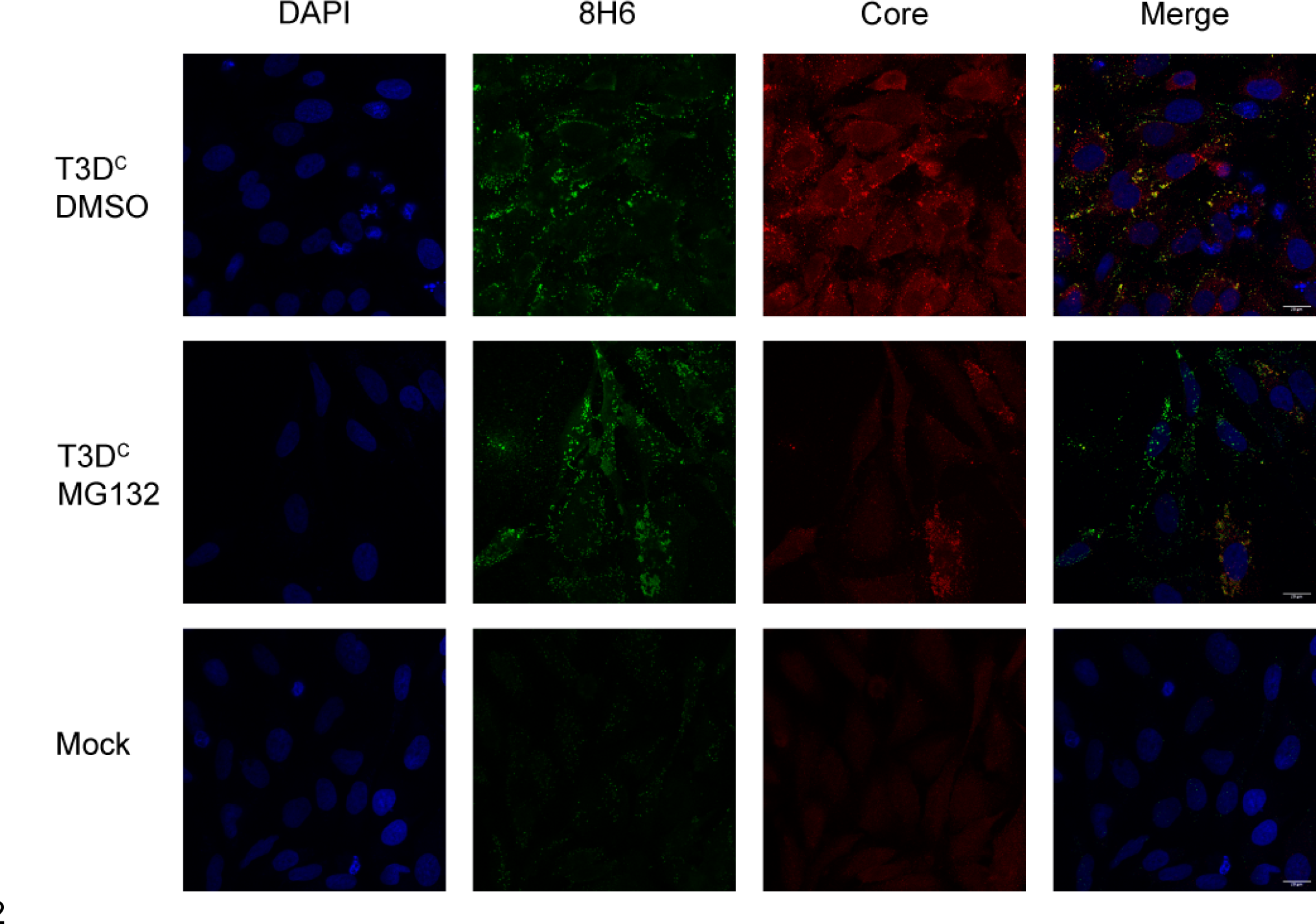
ISVPs fail to uncoat in presence of MG132. HeLa cells pretreated with CHX (100 μg/ml) and either DMSO or MG132 (10 μM) were infected with 10^5^ ISVPs of T3D^C^/cell. Cells were fixed 2 h post infection and immunostained using native μ1 specific 8H6 mAb (green), anti-core serum (red), and DAPI (blue). Uninfected cells were also stained to control for background staining.

One key step in the delivery of cores into the cytoplasm by ISVPs is the conversion to ISVP*s. ISVP-to-ISVP* transition results in a conformational change in the particle. The most dramatic conformational change is in the δ fragment and this conformational change allows the release of viral pore forming peptides for membrane penetration and core delivery. Since ISVPs were internalized but failed to deliver cores in the presence of MG132, we evaluated whether this conformational change was impacted by MG132. Conformational change in δ can detected by a conformationally sensitive mAb, 4A3, which detects δ in an altered ISVP* conformation but not the native, ISVP conformation (18). We observed that 4A3 staining was readily detected 2 h following infection with ISVPs in the presence of DMSO indicating that ISVPs have undergone ISVP-to-ISVP* conversion (Figure 5). Consistent with ISVP* conversion and matching the data in Figure 4 above, we observed that core staining was also visible in DMSO treated cells. In contrast, in the presence of MG132, minimal 4A3 or core staining was visible. These data indicate that inhibition of the proteasome impacts conformational changes accompanying ISVP-to-ISVP* transition.

**Figure 5.**
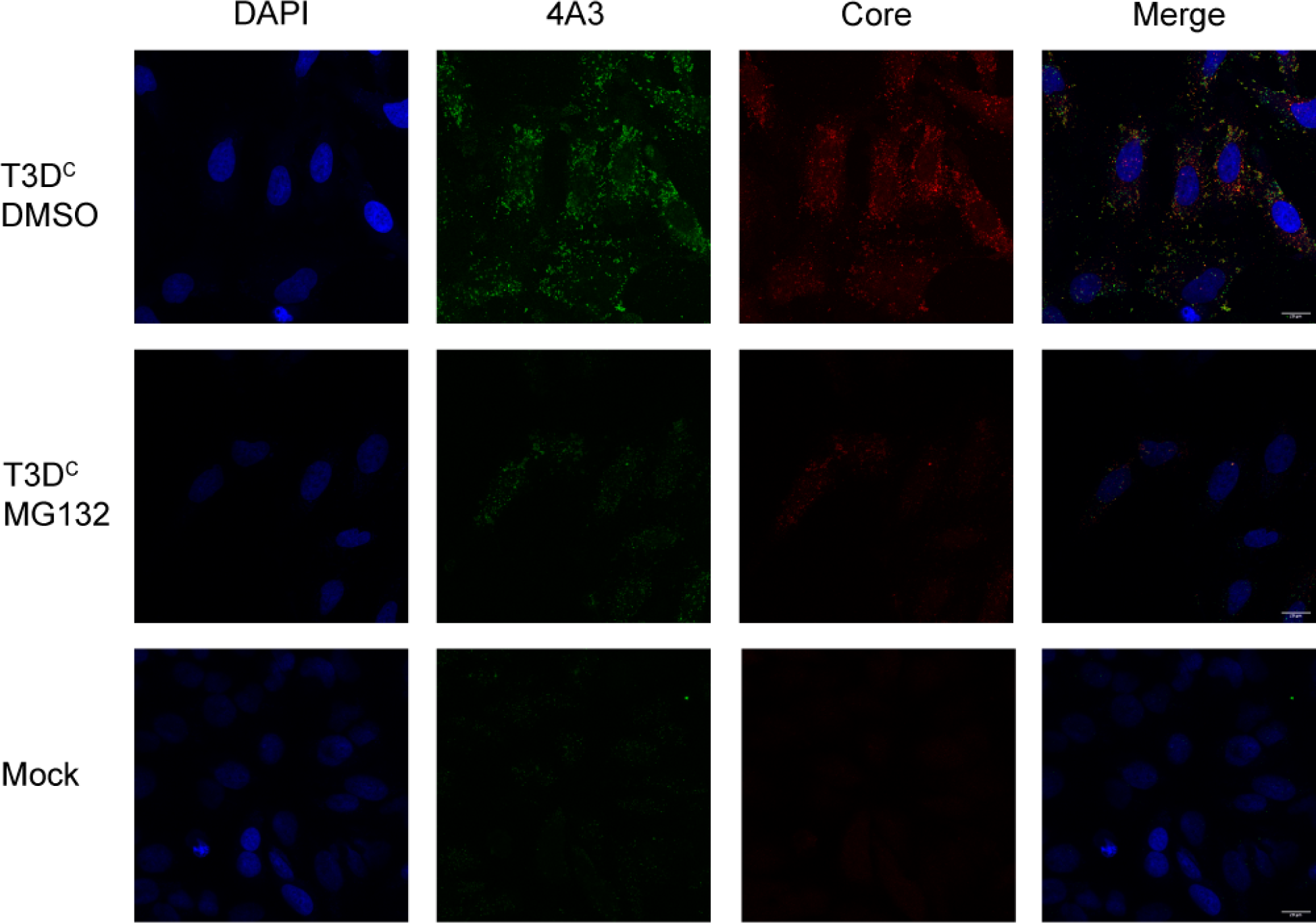
ISVPs fail to convert to ISVP*s in presence of MG132. HeLa cells pretreated with CHX (100 μg/ml) and either DMSO or MG132 (10 μM) were infected with 10^5^ ISVPs of T3D^C^/cell. Cells were fixed 2 h post infection and immunostained using altered μ1-specific 4A3 mAb (green), anti-core serum (red), and DAPI (blue). Uninfected cells were also stained to control for background staining.

ISVP-to-ISVP* conformational changes allow generation of additional μ1N fragments by autocleavage of μ1 (41). Further, these changes result in release of μ1N and φ from the particle (42, 43). These fragments facilitate penetration of the membrane and delivery of the cores into the cytoplasm (44, 45). Membrane penetration by ISVPs can be studied using the ribonucleotoxin, α-sarcin (18, 46, 47). α-sarcin does not affect protein synthesis of uninfected cells because it cannot cross the host membrane. However, it can be delivered into the cytoplasm in the wake of entering ISVPs and stop host protein synthesis. Using puromycin incorporation as a readout of protein synthesis (48), we observed that in comparison to uninfected cells, infection of DMSO treated cells with ISVPs allowed efficiently delivery of α-sarcin and strongly reduced the amount of puromycin that could be incoporated (Figure 6A). Though infection of MG132 treatment also blocked puromycin incorporation, this effect was less potent. Similar results were obtained at lower MOI of infection and with the T1L strain (data not shown). These data indicate that α-sarcin was less efficiently delivered into the cytoplasm in the presence of MG132. These data suggest that ISVPs fail to efficiently penetrate host membranes when proteasome activity is blocked.

**Figure 6.**
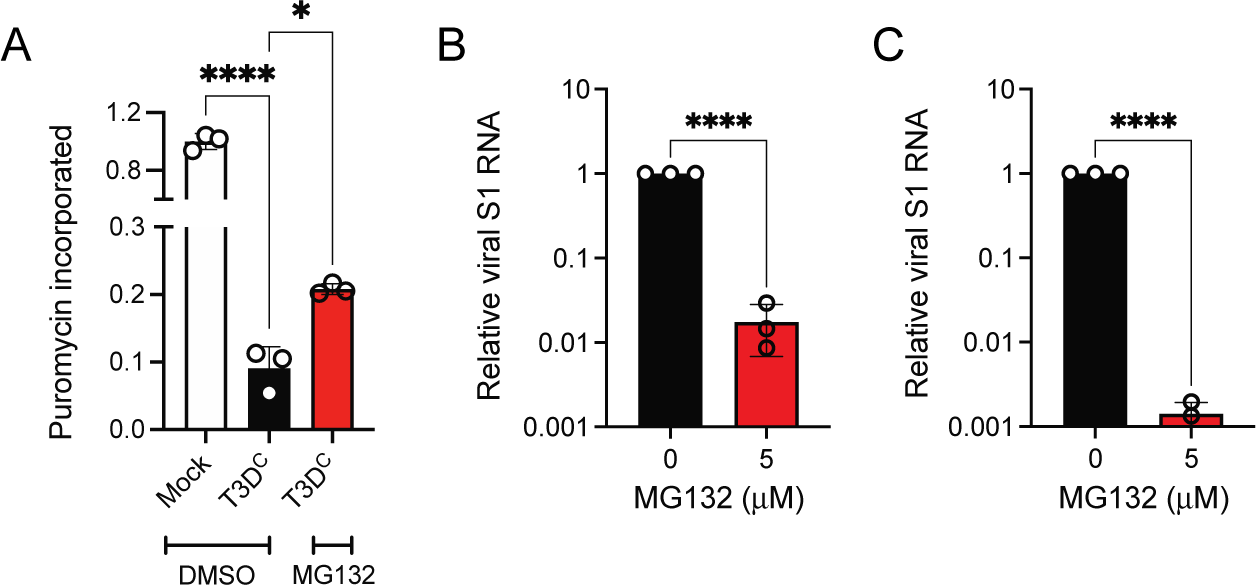
Uncoating defect in the presence of MG132 prevents membrane penetration and consequent viral gene expression. (A) HeLa cells pretreated with DMSO or MG132 (5 μM) were adsorbed with 10^5^ ISVPs of T3D^C^ per cell. Cells were infected for 2 h using DMSO or MG132 containing media supplemented with 50 μg/ml α-sarcin. 30 min prior to fixation, cells were incubated with 20 μg/ml puromycin. Puromycin incorporation was quantified by indirect immunofluorescence using a LI-COR Odyssey scanner. Protein synthesis efficiency was determined by calculating intensity ratios at 800 nm (green fluorescence) representing puromycin and 700 nm (red fluorescence) representing the cell monolayer. The 800/700 intensity ratio in uninfected cells was set to 1. Infectivity for each independent infection and the sample mean are shown. Error bars indicate SD. Error bars indicate SD. *, P < 0.05; ****, P < 0.001 as determined by one way ANOVA with Dunnett’s multiple comparison test. (B) HeLa cells were adsorbed with 10 PFU ISVPs of T3D^C^ per cell. Following incubation in the presence of DMSO or MG132 (5 μM) for 6 h, Levels of accumulated T3D S1 relative to GAPDH mRNA was quantified by qPCR and comparative CT analysis. The ratio of T3D S1 RNA to GAPDH in DMSO-treated cells was set to 1. Mean values for three independent infections are shown. Error bars indicate SD. P values were determined by student’s t-test. ****, p < 0.001. (C) L929 cells were adsorbed with 1 PFU ISVPs of T3D^C^ per cell. Following incubation in the presence of DMSO or MG132 (5 μM) for 6 h h, Levels of accumulated T3D S1 relative to GAPDH mRNA was quantified by qPCR and comparative CT analysis. The ratio of T3D S1 RNA to GAPDH in DMSO-treated cells was set to 1. Mean values for three independent infections are shown. Error bars indicate SD. P values were determined by student’s t-test. ****, P < 0.001.

Successful ISVP-to-ISVP* and membrane penetration allows delivery of the cores into the cytoplasm. Following removal of the altered δ fragment present on the cores, the particles begin transcribing viral RNA using particle-associated transcriptional machinery (49, 50). Based on reduced ISVP-to-ISVP* conversion and membrane penetration, we reasoned that viral mRNA transcription following core delivery will be reduced by MG132. To measure transcription from delivered cores, we measured the level of viral mRNA using RT-qPCR at 6 h following infection with ISVPs. We used the S1 gene transcript as a surrogate for other viral mRNAs. We found that in comparison to mRNA accumulated in cells in the presence of DMSO, there was significantly (100-fold) lower level of S1 mRNA in cells treated with MG132 (Figure 6B). These data provide further support for the idea that MG132 diminishes reovirus infection by decreasing events in entry following ISVP formation. Using transcription from incoming cores as a readout, we determined that the dependence of ISVP-to-ISVP* on proteasome activity extends to other cell types. The accumulation of S1 RNA at 6 h following infection with ISVPs also was significantly lower in L929 cells treated with MG132 (Figure 6C). Together our results indicate that proteasome activity dependent uncoating of reovirus is required for successful infection.

## DISCUSSION

In this paper, we evaluated the effect of proteasome activity on reovirus infection. Our results reveal that proteasome inhibitors affect reovirus infection in two ways. First, they reduce conversion of virion to ISVP by inhibiting cathepsin activity. Second, proteasome inhibitors impact ISVP uncoating steps associated with delivery of cores into the cytoplasm thereby preventing the successful initiation of the reovirus replication cycle.

Among these effects, it is known that MG132 and other proteasome inhibitors inhibit the activity of proteases other than those that reside within the proteasome (33). Therefore, their impact on cathepsin activity and consequent effect on reovirus disassembly isn’t unexpected. However, our results should serve as a caution to those studying the effect of proteasome on turnover of host and viral proteins in reovirus infected cells. Alternate tools such as inhibition of individual or groups of E3 ubiquitin ligases may have to be used as tools for such experiments.

The results identifying a role of proteasome activity in uncoating of already disassembled ISVPs is more interesting because this step is poorly understood. On ISVPs, the μ1 protein is present as two particle associated fragments, μ1δ and φ (3, 4, 40). The conformation of μ1 on the particle resembles that which is present on the native virion, even though its binding partner, σ3 is absent. ISVPs undergo conformational changes to convert to ISVP*s (18). During this change, μ1δ is further cleaved to form μ1N and δ (41). Conformational changes in the particle result in rearrangement of δ and concomitant release of μ1N, φ and σ1 (43). Our evidence suggests that inhibition of the proteasome activity inhibits this step. Interestingly, proteasomes are present in the cytoplasm and ISVP-to-ISVP* conversion occurs in an early uptake vesicle or in an endosome. Thus, it seems unlikely that a capsid associated viral protein is being targeted by the proteasome to mediate uncoating.

Consistent with this, in purified virus preparations used to initiate infection in this and many other studies, capsid proteins migrate at their expected molecular weight in SDS-PAGE gels and are not thought to be ubiquitin modified. Immunoblotting purified virus preparations with ubiquitin-specific antibodies also supports this idea (data not shown). We also did not find detectable changes to the ubiquitination status of the δ fragment of μ1 from internalized virions (data not shown). Thus, this effect of proteasome inhibition on virus uncoating seems distinct from that described for vaccinia virus and African swine fever virus (51–53), where specific viral proteins are targeted for degradation by the proteasome in the cytoplasm.

Based on studies with Kaposi Sarcoma-associated Herpesvirus and murine hepatitis virus, it is suggested that proteasome inhibition prevents particle escape or genome delivery from the endosome by affecting endosome maturation (54, 55). In most cell types, reovirus disassembly and uncoating occurs in the endosomal compartment and transit through late endosomes is required for cell entry (37, 49, 56, 57). Indeed, perturbing maturation of endosomes using pharmacologic or genetic manipulation blocks reovirus entry. While inhibition of infection initiated by reovirus particles may be similarly affected by effects on endosomal maturation, we believe that our observation that infection initiated by ISVPs is also inhibited by proteasome inhibitors indicates a distinct mechanism. When infection is initiated by in vitro generated ISVPs, ISVPs do not require uptake pathways that deliver particles to the endosomes (11, 58). Further, treatments that inhibit endosomal maturation or affect endosomal protease activity do not inhibit infection initiated by ISVPs (32, 37, 49, 56, 57, 59). We therefore think that proteasome inhibition affects infection by ISVPs by a previously unknown mechanism. It should be noted that while in vitro generated ISVPs are tools to investigate viral entry, such particles are formed in the lumen of the intestinal and respiratory tract (60–62). These in vivo generated ISVPs are thought to have entry requirements that match those of in vitro generated ISVPs.

Exactly what triggers ISVP-to-ISVP* conformational changes in the particle is unknown but our model posits that μ1 fragments cooperate with host membranes to promote this transition (63, 64). No cellular proteins are known to participate in this process. We propose that by altering the levels of cellular proteins involved in lipid metabolism, proteasome inhibitors alter the lipid composition, and that this results in defects in uncoating of ISVPs.

## FIGURE LEGENDS

## REFERENCES

1. Yamauchi Y, Helenius A. 2013. Virus entry at a glance. J Cell Sci 126:1289–95.

2. Dermody TS, Parker JC, Sherry B. 2013. Orthoreoviruses, p 1304–1346. In Knipe DM, Howley PM (ed), Fields Virology, Sixth ed, vol 2. Lippincott Williams & Wilkins, Philadelphia.

3. Zhang X, Ji Y, Zhang L, Harrison SC, Marinescu DC, Nibert ML, Baker TS. 2005. Features of reovirus outer capsid protein m1 revealed by electron cryomicroscopy and image reconstruction of the virion at 7.0 Å resolution. Structure 13:1545–1557.

4. Dryden KA, Wang G, Yeager M, Nibert ML, Coombs KM, Furlong DB, Fields BN, Baker TS. 1993. Early steps in reovirus infection are associated with dramatic changes in supramolecular structure and protein conformation: analysis of virions and subviral particles by cryoelectron microscopy and image reconstruction. Journal of Cell Biology 122:1023–1041.

5. Danthi P, Holm G. H., Stehle T., and Dermody T.S. . 2013. Reovirus receptors, cell entry, and signaling. *In* Pöhlmann S, and Simmons G. (ed), Viral Entry into Cells, Georgetown, TX.

6. Barton ES, Connolly JL, Forrest JC, Chappell JD, Dermody TS. 2001. Utilization of sialic acid as a coreceptor enhances reovirus attachment by multistep adhesion strengthening. J Biol Chem 276:2200–11.

7. Barton ES, Forrest JC, Connolly JL, Chappell JD, Liu Y, Schnell FJ, Nusrat A, Parkos CA, Dermody TS. 2001. Junction adhesion molecule is a receptor for reovirus. Cell 104:441–51.

8. Kirchner E, Guglielmi KM, Strauss HM, Dermody TS, Stehle T. 2008. Structure of reovirus sigma1 in complex with its receptor junctional adhesion molecule-A. PLoS Pathog 4:e1000235.

9. Reiss K, Stencel JE, Liu Y, Blaum BS, Reiter DM, Feizi T, Dermody TS, Stehle T. 2012. The GM2 glycan serves as a functional coreceptor for serotype 1 reovirus. PLoS Pathog 8:e1003078.

10. Reiter DM, Frierson JM, Halvorson EE, Kobayashi T, Dermody TS, Stehle T. 2011. Crystal structure of reovirus attachment protein sigma1 in complex with sialylated oligosaccharides. PLoS Pathog 7:e1002166.

11. Borsa J, Morash BD, Sargent MD, Copps TP, Lievaart PA, Szekely JG. 1979. Two modes of entry of reovirus particles into L cells. Journal of General Virology 45:161–170.

12. Maginnis MS, Forrest JC, Kopecky-Bromberg SA, Dickeson SK, Santoro SA, Zutter MM, Nemerow GR, Bergelson JM, Dermody TS. 2006. Beta1 integrin mediates internalization of mammalian reovirus. J Virol 80:2760–70.

13. Maginnis MS, Mainou BA, Derdowski A, Johnson EM, Zent R, Dermody TS. 2008. NPXY motifs in the beta1 integrin cytoplasmic tail are required for functional reovirus entry. J Virol 82:3181–91.

14. Aravamudhan P, Raghunathan K, Konopka-Anstadt J, Pathak A, Sutherland DM, Carter BD, Dermody TS. 2020. Reovirus uses macropinocytosis-mediated entry and fast axonal transport to infect neurons. PLoS Pathog 16:e1008380.

15. Schulz WL, Haj AK, Schiff LA. 2012. Reovirus uses multiple endocytic pathways for cell entry. J Virol 86:12665–75.

16. Koehler M, Petitjean SJL, Yang J, Aravamudhan P, Somoulay X, Lo Giudice C, Poncin MA, Dumitru AC, Dermody TS, Alsteens D. 2021. Reovirus directly engages integrin to recruit clathrin for entry into host cells. Nat Commun 12:2149.

17. Ebert DH, Deussing J, Peters C, Dermody TS. 2002. Cathepsin L and cathepsin B mediate reovirus disassembly in murine fibroblast cells. J Biol Chem 277:24609–17.

18. Chandran K, Parker JS, Ehrlich M, Kirchhausen T, Nibert ML. 2003. The delta region of outer-capsid protein m1 undergoes conformational change and release from reovirus particles during cell entry. Journal of Virology 77:13361–13375.

19. Chandran K, Farsetta DL, Nibert ML. 2002. Strategy for nonenveloped virus entry: a hydrophobic conformer of the reovirus membrane penetration protein m1 mediates membrane disruption. Journal of Virology 76:9920–9933.

20. Kniert J, Dos Santos T, Eaton HE, Jung Cho W, Plummer G, Shmulevitz M. 2022. Reovirus uses temporospatial compartmentalization to orchestrate core versus outercapsid assembly. PLoS Pathog 18:e1010641.

21. Virgin HW, IV, Mann MA, Fields BN, Tyler KL. 1991. Monoclonal antibodies to reovirus reveal structure/function relationships between capsid proteins and genetics of susceptibility to antibody action. Journal of Virology 65:6772–6781.

22. Kobayashi T, Ooms LS, Ikizler M, Chappell JD, Dermody TS. 2010. An improved reverse genetics system for mammalian orthoreoviruses. Virology 398:194–200.

23. Berard A, Coombs KM. 2009. Mammalian reoviruses: propagation, quantification, and storage. Curr Protoc Microbiol Chapter 15:Unit15C 1.

24. Nibert ML, Chappell JD, Dermody TS. 1995. Infectious subvirion particles of reovirus type 3 Dearing exhibit a loss in infectivity and contain a cleaved sigma 1 protein. J Virol 69:5057–67.

25. Middleton JK, Severson TF, Chandran K, Gillian AL, Yin J, Nibert ML. 2002. Thermostability of reovirus disassembly intermediates (ISVPs) correlates with genetic, biochemical, and thermodynamic properties of major surface protein mu1. J Virol 76:1051–61.

26. Iskarpatyoti JA, Willis JZ, Guan J, Morse EA, Ikizler M, Wetzel JD, Dermody TS, Contractor N. 2012. A rapid, automated approach for quantitation of rotavirus and reovirus infectivity. J Virol Methods 184:1–7.

27. Schmittgen TD, Livak KJ. 2008. Analyzing real-time PCR data by the comparative C(T) method. Nat Protoc 3:1101–8.

28. Rasband WS. Image J. U S National Institutes of Health, Bethesda, Maryland, USA:1997-2007.

29. McNamara AJ, Danthi P. 2020. Loss of IKK Subunits Limits NF-kappaB Signaling in Reovirus-Infected Cells. J Virol 94.

30. Lee DH, Goldberg AL. 1998. Proteasome inhibitors: valuable new tools for cell biologists. Trends Cell Biol 8:397–403.

31. Varfolomeev E, Blankenship JW, Wayson SM, Fedorova AV, Kayagaki N, Garg P, Zobel K, Dynek JN, Elliott LO, Wallweber HJ, Flygare JA, Fairbrother WJ, Deshayes K, Dixit VM, Vucic D. 2007. IAP antagonists induce autoubiquitination of c-IAPs, NF-kappaB activation, and TNFalpha-dependent apoptosis. Cell 131:669–81.

32. Sturzenbecker LJ, Nibert ML, Furlong DB, Fields BN. 1987. Intracellular digestion of reovirus particles requires a low pH and is an essential step in the viral infectious cycle. Journal of Virology 61:2351–2361.

33. Kisselev AF, Goldberg AL. 2001. Proteasome inhibitors: from research tools to drug candidates. Chem Biol 8:739–58.

34. Snyder AJ, Wang JC, Danthi P. 2018. Components of the reovirus capsid differentially contribute to stability. J Virol doi:10.1128/JVI.01894-18.

35. Abad AT, Danthi P. 2022. Early Events in Reovirus Infection Influence Induction of Innate Immune Response. J Virol 96:e0091722.

36. Bokiej M, Ogden KM, Ikizler M, Reiter DM, Stehle T, Dermody TS. 2012. Optimum length and flexibility of reovirus attachment protein sigma1 are required for efficient viral infection. J Virol 86:10270–80.

37. Mainou BA, Zamora PF, Ashbrook AW, Dorset DC, Kim KS, Dermody TS. 2013. Reovirus cell entry requires functional microtubules. MBio 4.

38. Reinisch KM, Nibert ML, Harrison SC. 2000. Structure of the reovirus core at 3.6 Å resolution. Nature 404:960–967.

39. Jiang J, Coombs KM. 2005. Infectious entry of reovirus cores into mammalian cells enhanced by transfection. J Virol Methods 128:88–92.

40. Pan M, Alvarez-Cabrera AL, Kang JS, Wang L, Fan C, Zhou ZH. 2021. Asymmetric reconstruction of mammalian reovirus reveals interactions among RNA, transcriptional factor micro2 and capsid proteins. Nat Commun 12:4176.

41. Nibert ML, Odegard AL, Agosto MA, Chandran K, Schiff LA. 2005. Putative autocleavage of reovirus m1 protein in concert with outer-capsid disassembly and activation for membrane permeabilization. Journal of Molecular Biology 345:461–474.

42. Agosto MA, Myers KS, Ivanovic T, Nibert ML. 2008. A positive-feedback mechanism promotes reovirus particle conversion to the intermediate associated with membrane penetration. Proc Natl Acad Sci U S A 105:10571–6.

43. Ivanovic T, Agosto MA, Zhang L, Chandran K, Harrison SC, Nibert ML. 2008. Peptides released from reovirus outer capsid form membrane pores that recruit virus particles. Embo J 27:1289–1298.

44. Odegard AL, Chandran K, Zhang X, Parker JS, Baker TS, Nibert ML. 2004. Putative autocleavage of outer capsid protein m1, allowing release of myristoylated peptide m1N during particle uncoating, is critical for cell entry by reovirus. Journal of Virology 78:8732–8745.

45. Snyder AJ, Danthi P. 2018. Cleavage of the C-Terminal Fragment of Reovirus mu1 Is Required for Optimal Infectivity. J Virol 92.

46. Danthi P, Kobayashi T, Holm GH, Hansberger MW, Abel TW, Dermody TS. 2008. Reovirus apoptosis and virulence are regulated by host cell membrane penetration efficiency. J Virol 82:161–72.

47. Martinez CG, Guinea R, Benavente J, Carrasco L. 1996. The entry of reovirus into L cells is dependent on vacuolar proton-ATPase activity. Journal of Virology 70:576–579.

48. Starck SR, Green HM, Alberola-Ila J, Roberts RW. 2004. A general approach to detect protein expression in vivo using fluorescent puromycin conjugates. Chem Biol 11:999–1008.

49. Snyder AJ, Abad AT, Danthi P. 2022. A CRISPR-Cas9 screen reveals a role for WD repeat-containing protein 81 (WDR81) in the entry of late penetrating viruses. PLoS Pathog 18:e1010398.

50. Ortega-Gonzalez P, Taylor G, Jangra RK, Tenorio R, de Castro IF, Mainou BA, Orchard RC, Wilen CB, Brigleb PH, Sojati J, Chandran K, Risco C, Dermody TS. 2021. Reovirus infection is regulated by NPC1 and endosomal cholesterol homeostasis. bioRxiv.

51. Mercer J, Snijder B, Sacher R, Burkard C, Bleck CK, Stahlberg H, Pelkmans L, Helenius A. 2012. RNAi screening reveals proteasome- and Cullin3-dependent stages in vaccinia virus infection. Cell Rep 2:1036–47.

52. Schmidt FI, Bleck CK, Reh L, Novy K, Wollscheid B, Helenius A, Stahlberg H, Mercer J. 2013. Vaccinia virus entry is followed by core activation and proteasome-mediated release of the immunomodulatory effector VH1 from lateral bodies. Cell Rep 4:464–76.

53. Barrado-Gil L, Galindo I, Martinez-Alonso D, Viedma S, Alonso C. 2017. The ubiquitin-proteasome system is required for African swine fever replication. PLoS One 12:e0189741.

54. Greene W, Zhang W, He M, Witt C, Ye F, Gao SJ. 2012. The ubiquitin/proteasome system mediates entry and endosomal trafficking of Kaposi’s sarcoma-associated herpesvirus in endothelial cells. PLoS Pathog 8:e1002703.

55. Yu GY, Lai MM. 2005. The ubiquitin-proteasome system facilitates the transfer of murine coronavirus from endosome to cytoplasm during virus entry. J Virol 79:644–8.

56. Mainou BA, Dermody TS. 2011. Src kinase mediates productive endocytic sorting of reovirus during cell entry. J Virol 85:3203–13.

57. Mainou BA, Dermody TS. 2012. Transport to late endosomes is required for efficient reovirus infection. J Virol 86:8346–58.

58. Boulant S, Stanifer M, Kural C, Cureton DK, Massol R, Nibert ML, Kirchhausen T. 2013. Similar uptake but different trafficking and escape routes of reovirus virions and infectious subvirion particles imaged in polarized Madin-Darby canine kidney cells. Mol Biol Cell 24:1196–207.

59. Ebert DH, Wetzel JD, Brumbaugh DE, Chance SR, Stobie LE, Baer GS, Dermody TS. 2001. Adaptation of reovirus to growth in the presence of protease inhibitor E64 segregates with a mutation in the carboxy terminus of viral outer-capsid protein sigma3. J Virol 75:3197–206.

60. Bodkin DK, Nibert ML, Fields BN. 1989. Proteolytic digestion of reovirus in the intestinal lumens of neonatal mice. Journal of Virology 63:4676–4681.

61. Bass DM, Bodkin D, Dambrauskas R, Trier JS, Fields BN, Wolf JL. 1990. Intraluminal proteolytic activation plays an important role in replication of type 1 reovirus in the intestines of neonatal mice. Journal of Virology 64:1830–1833.

62. Nygaard RM, Golden JW, Schiff LA. 2012. Impact of host proteases on reovirus infection in the respiratory tract. Journal of virology 86:1238–43.

63. Snyder AJ, Danthi P. 2015. Lipid Membranes Facilitate Conformational Changes Required for Reovirus Cell Entry. J Virol 90:2628–38.

64. Snyder AJ, Danthi P. 2016. Lipids Cooperate with the Reovirus Membrane Penetration Peptide to Facilitate Particle Uncoating. J Biol Chem 291:26773–26785.

